# Distinct neural architectures prevent off-task thought in different task contexts

**DOI:** 10.64898/2026.04.23.719614

**Authors:** Chen Chen, Katya Krieger-Redwood, Meichao Zhang, Lidón Marin-Marin, Ximing Shao, Jonathan Smallwood, Elizabeth Jefferies

## Abstract

Prevailing models propose that sustained focus is a product of top-down control that amplifies task-relevant representations while suppressing distraction. We challenge this view, demonstrating that the neural signatures of mental focus differ across cognitive modes. In two neuroimaging studies, we used a paradigm that creates conflict between external task goals and self-generated thought and modelled neural activity as a function of trial-by-trial subjective focus to identify how people stay on task in contexts varying in memory engagement. In Study 1, on-task thoughts were associated with distinct neural architectures in arithmetic and comprehension: arithmetic-related focus recruited visual–motor regions, while comprehension-related focus was characterized by the recruitment of memory systems and reduced activation of control regions. Study 2 extended these findings, with both externally directed comprehension and internal memory retrieval associated with a shared neural profile of reduced control engagement, despite reliance on distinct memory systems.

Furthermore, focus covaried with activity in heteromodal semantic regions across both reading and listening, suggesting semantic engagement is a primary marker of being on-task during comprehension. Our findings indicate that sustained focus during memory-guided cognition is characterized by mutual disengagement between control and memory networks. This discovery challenges the view that top-down supervision is a universal requirement for task focus, since sustained attention is associated with distinct neural configurations in different cognitive modes.

## Introduction

Task focus is conventionally viewed as a product of deliberate, top-down control ^1, 2, 3^. Yet, the brain can sustain remarkable focus during immersive memory recall with minimal apparent effort, challenging the assumption that task focus always depends on executive regulation. Prior work demonstrates that task engagement sharpens neural responses in relevant cortical regions and suppresses competing information by modulating the connectivity between control, sensory, and default mode networks (DMN; e.g., ref. ^4, 5, 6^). However, this account cannot explain forms of sustained cognition that unfold independently of control networks. Here, we test whether the brain can maintain thoughts focused on performing a task through the self-organizing dynamics of memory systems, without continuous executive oversight.

The fronto-parietal control network (FPCN) maintains goal-relevant representations and typically responds more strongly when demands increase ^7, 8, 9^. Through its subregions, FPCN aligns functionally with either DMN or the dorsal attention network, thereby supporting internal and external modes of cognition ^10, 11, 12, 13, 14, 15^. A classic executive-failure hypothesis proposes that mind-wandering during externally oriented tasks such as reading reflects a breakdown in top-down control. When frontal–parietal control systems fail to prioritize task-relevant information, there is insufficient suppression of internally generated content within the DMN ^16, 17^. Although earlier accounts often focused on mind-wandering, subjective mental states form a continuum ranging from on-task states, in which ongoing cognition is aligned with task demands, to off-task states, in which thought becomes decoupled from the current goal ^18, 19^. Under this executive-control account, moments of greater subjective focus might therefore be expected to coincide with stronger control engagement.

The DMN, conversely, is classically assumed to deactivate in attention-demanding contexts ^20, 21^, and to activate during unconstrained, self-generated thought ^22, 23^. Yet a growing literature shows that DMN activity can accompany goal-directed cognition when memory retrieval or semantic comprehension is required ^24, 25^. Some cognitive states might also be sustained via memory-based systems: dorsomedial and core DMN subsystems are recruited during both text comprehension and autobiographical recall respectively, with stronger core DMN activation observed when participants are more focused on memory recall than comprehension ^25, 26^. These patterns indicate that semantic engagement might be a common mechanism supporting focus across internally and externally oriented cognition ^27^, and that memory networks are positively associated with task focus in some cognitive modes. Established theories of comprehension also treat meaning-construction as memory-based: text is understood by activating and integrating knowledge in long-term memory ^28, 29^, while naturalistic neuroimaging treats default mode engagement as a marker of meaning-construction over extended, ecologically valid input ^30, 31, 32^. These frameworks characterise how meaning is constructed from input; however, they do not address what sustains on-task states supporting comprehension, and whether these states involve the engagement of control processes alongside meaning.

Behavioral evidence has already started to show that the mechanisms that sustain on-task thought may be different in inherently semantic task contexts, such as reading, compared with non-semantic tasks. In externally oriented attention tasks, increasing task demands are often associated with less mind-wandering ^33, 34, 35, 36^. This pattern is consistent with executive-control accounts, according to which greater control engagement under higher demands may help maintain the task goal and suppress task-unrelated thought. However, studies of reading comprehension reveal the opposite: people report more off-task thought when texts are difficult ^37, 38, 39, 40^. One possibility is that successful semantic construction itself contributes to the maintenance of focus during comprehension. Yet little is known about the neurobiological processes that can explain this dissociation in subjective task focus.

To test these ideas, we conducted two functional MRI studies examining how large-scale brain networks support sustained focus across internal and external cognitive modes. In Study 1, participants performed externally oriented tasks -- arithmetic or sentence comprehension -- with each trial preceded by autobiographical memory cues, creating competition between task goals and self-generated thought. Trial-by-trial focus ratings allowed us to model how fluctuations in attention shape neural activity and network coupling. Study 2 extended and replicated these effects by comparing sentence comprehension and autobiographical memory retrieval tasks, presented either alone or in competition, across spoken and written modalities. This design enabled us to test whether similar mechanisms are associated with focus across semantic and memory-based states, and whether modality influences these processes. Critically, our approach differs from prior neuroimaging studies of comprehension, which model neural activity locked to properties of the stimulus, such as text difficulty, coherence or narrative structure ^31, 41, 42^. Here we instead model activity that tracks participants’ trial-by-trial subjective focus while holding the task and stimuli constant, isolating the neural signature of being on task from that of the task itself.

Across analyses, we observed distinct neural pathways associated with focus depending on cognitive context. In Study 1, focus in arithmetic involved reduced engagement of memory regions, whereas comprehension-related focus involved deactivation of control regions. Study 2 showed that both comprehension and autobiographical memory retrieval were associated with greater reductions in control engagement at higher levels of focus, with comprehension additionally associated with reduced activity in core DMN regions. Heteromodal semantic regions were associated with higher focus across reading and listening tasks, indicating that memory rather than sensory systems covary with sustained comprehension. Conjunction analyses across studies 1 and 2 confirmed consistent semantic activation and control deactivation during focused comprehension. Finally, bidirectional connectivity analyses revealed mutual decoupling between control and semantic regions during focused comprehension, consistent with a pattern in which memory-based systems coincide with more stable cognitive states in the absence of executive engagement. Together, these findings challenge traditional executive-control accounts, indicating that in some cognitive modes, sustained thought reflects the engagement of memory architectures rather than top-down supervision.

## Results

### Study 1: Differential focus effects during semantic and non-semantic tasks

Study 1 examined how task focus is sustained across semantic and non-semantic contexts to test the hypothesis that these distinct modes of cognition are sustained in different ways. In a within-subjects design (N=34), participants performed comprehension (semantic) and arithmetic (non-semantic) tasks that varied in difficulty (Easy vs. Hard; see Materials and Methods). Prior to scanning, participants generated personal thoughts linked to cue phrases (e.g., “Lottery win”) to induce internally oriented, memory-based distraction. These cues were designed to trigger autobiographical retrieval during fMRI while participants either read factual sentences or solved arithmetic problems (serial addition). After each trial, participants rated their task focus on a 1–7 scale (1 = thinking about other things, attention fully diverted from task; 7 = thoughts fully on task; Fig. 1A).

**Fig 1.**
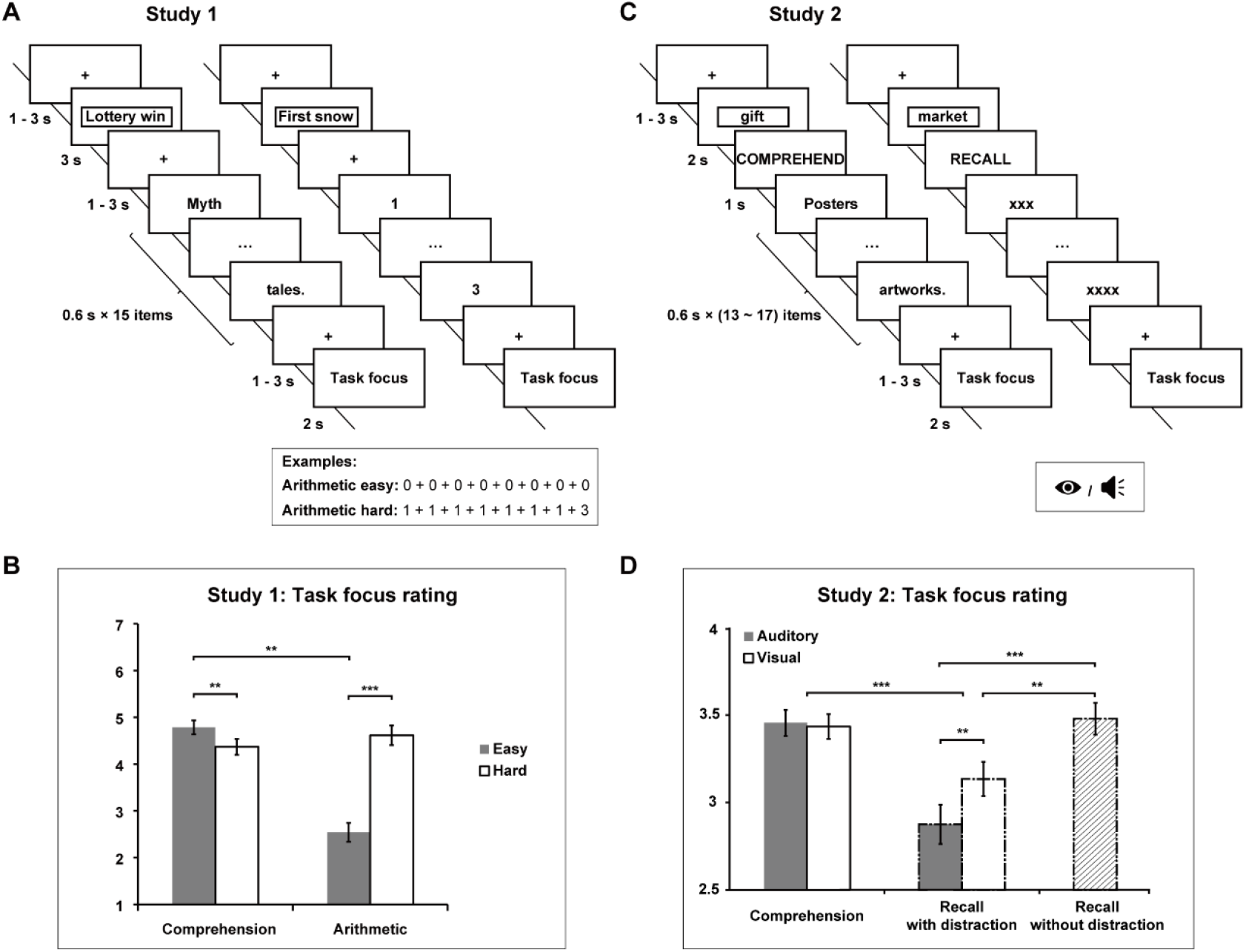
Task illustration and task focus ratings in two studies. (A) Task procedure in study 1. Participants retrieved personal thoughts linked to an autobiographical cue phrase (phrase with a box, e.g., “Lottery win”), then performed a comprehension or arithmetic task. At the end of each trial, participants rated their level of focus on a scale from 1 to 7. (B) Ratings of task focus in study 1. The color of solid bars indicates task difficulty (grey, easy; white, hard). Error bars represent the standard error of the mean (SE). (C) Task procedure in study 2. Participants recalled personal memories from a cue word (word with a box, e.g., “gift”) in the scanner, and then performed either a comprehension or recall task. In the comprehension task, participants read or listened to factual sentences. In the recall task, participants recalled personal memories while ignoring visual or auditory sentences, or without distraction. At the end of each trial, participants rated their level of focus on the intended task on a scale from 1 to 4. (D) Ratings of task focus in study 2. The color of solid bars represents the sentence modality (grey, auditory; white, visual). The grid bar represents focus during recall without distraction. Error bars represent the standard error of the mean (SE). \*\*\**p* < 0.001, \*\**p* < 0.01, Bonferroni corrected.

#### Behavioral results

A repeated-measures ANOVA examining task focus ratings explored the effects of task (comprehension/arithmetic) and difficulty (easy/hard). This revealed main effects of task (*F*(1, 33) = 54.08, *p* < .001, *η*^*2*^ = .62), difficulty (*F*(1, 33) = 27.47, *p* < .001, *η*^*2*^ = .45), and a task-by-difficulty interaction (*F*(1, 33) = 58.60, *p* < .001, *η*^*2*^ = .64) (Fig. 1B). Post-hoc Bonferroni-corrected *t*-tests showed that participants were more focused on easy than hard comprehension (*t*(33) = 3.348, Bonferroni-corrected *p* = .002), and more focused on hard than easy arithmetic (*t*(33) = −7.01, Bonferroni-corrected *p* < .001).

This behavioral dissociation suggests that the mechanisms sustaining task focus differ across domains. In the arithmetic task, participants achieve higher focus during harder trials by actively suppressing mind-wandering to meet task demands. In contrast, comprehension relies on engagement of the semantic system: focus is higher in easier texts, where visual-semantic processing more effectively stabilizes attention on the material.

#### Parametric effects of task focus in comprehension and arithmetic

To identify neural responses associated with subjective fluctuations in focus, we implemented a univariate general linear model (GLM; see Materials and Methods). Task difficulty was modelled categorically, while trial-by-trial ratings of task focus were entered as a parametric regressor across all trials. This approach isolated neural activity that covaried with moment-to-moment variation in focus, independent of mean activation differences between easy and hard conditions (see below; Fig. 2). Main effects of task are shown in SI Appendix, Fig. S1. Supplementary analyses confirmed highly similar effects of focus in the hard and easy trials of both tasks when these were analyzed separately, and no significant differences in focus effects between hard and easy trials for either task (SI Appendix, Fig. S2).

**Fig 2.**
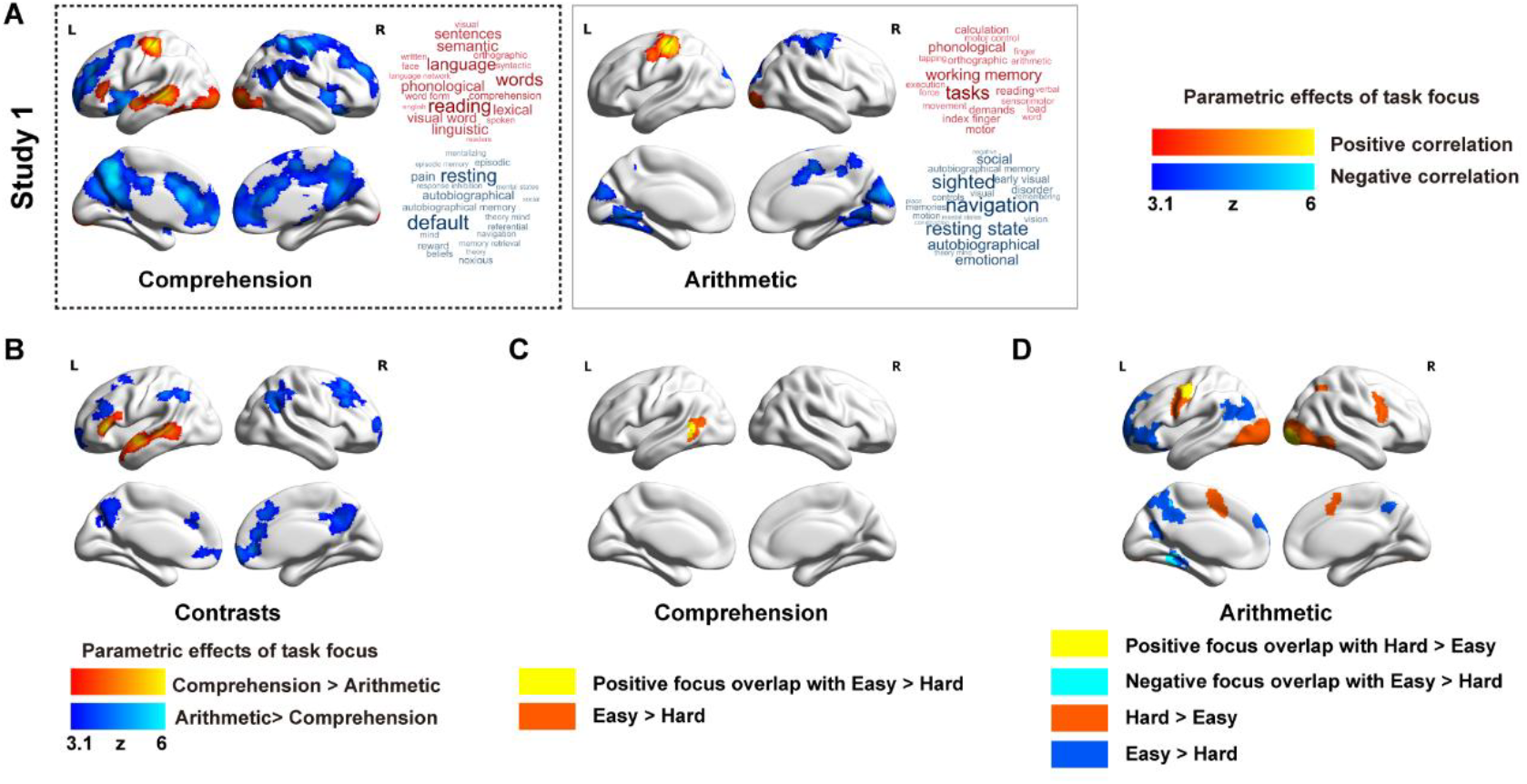
Parametric effects of task focus in Study 1. (A) These brain regions showed greater activation (red) or greater deactivation (blue) when participants reported better focus on the comprehension or arithmetic task. The word clouds indicate cognitive decoding of each map using Neurosynth: the font size of each item illustrates its importance, and the color indicates its association (red = positive correlation, blue = negative correlation). (B) A comparison of regions showing greater focus effect during comprehension (red) or arithmetic task (blue). (C) Overlap between focus effect and task difficulty during comprehension. Red indicates regions showing greater activation during easy compared to hard comprehension. Yellow indicates regions associated with increased focus overlap with those more active during easy trials. (D) Overlap between focus effect and task difficulty during arithmetic. Red indicates regions showing greater activation during hard compared to easy arithmetic; dark blue indicates the opposite. Yellow indicates regions associated with increased focus overlap with those more active during hard trials, while bright blue indicates regions associated with decrease focus overlap with those more active during easy trials. Source data of unthresholded focus effect maps in A-B are available at NeuroVault (https://neurovault.org/collections/UXFXQCND/).

To examine how focus-related neural patterns map onto large-scale brain networks, we assessed the voxel-level overlap between thresholded focus activation maps and the 17- network parcellation defined by ref. ^43^. Networks were split by hemisphere to capture potential lateralization effects, given the left-lateralization of semantic cognition. Results are shown in SI Appendix, Fig. S3, with separate bars for each hemisphere and colors indicating different networks. To simplify visualization, networks with fewer than 100 overlapping voxels are grouped as “other” (all networks are shown in OSF; https://osf.io/cqbhv/overview?view_only=ea533ec03ad3471a8336cc32107c15f6).

Greater focus during comprehension was associated with increased activation in the bilateral occipital cortex, left precentral/postcentral gyrus, inferior frontal gyrus (IFG), middle temporal gyrus (MTG), superior temporal gyrus (STG), and inferior temporal regions; see Fig. 2A. Using Neurosynth ^44^, we performed meta-analysis to reveal functional associations with these regions. Cognitive decoding of the focus map for comprehension identified terms such as “reading”, “words”, and “language”. These regions fell largely within visual, somatomotor, auditory and left-lateralized dorsomedial DMN networks, as defined by the ref. ^43^ parcellation of intrinsic connectivity (see SI Appendix, Fig. S3).

Higher focus during comprehension also elicited greater deactivation in the bilateral anterior and posterior cingulate, precuneus, insula, frontal pole, superior frontal gyrus (SFG), middle frontal gyrus (MFG), and right lateral parietal cortex; Fig. 2A. Cognitive decoding of this map identified terms associated with “default”, “resting”, and “autobiographical” as well as “pain”, “noxious” and “response inhibition”. These regions largely fell within bilateral core DMN, control, salience and dorsal attention networks (SI Appendix, Fig. S3).

During the arithmetic task, higher focus was associated with increased activation in left precentral/postcentral gyrus and right occipital pole; cognitive decoding identified terms such as “tasks”, “working memory”, and “phonological” (Fig. 2A). These areas fell in the left somatomotor and right visual network (SI Appendix, Fig. S3). Higher focus was also associated with deactivation in the bilateral cuneal cortex, lingual gyrus, fusiform gyrus, precuneus, and right cingulate, precentral/postcentral gyrus, and associated with terms like “navigation”, “sighted”, “resting” and “autobiographical”. These areas of deactivation were largely within bilateral visual network and right SMN.

This analysis confirms distinct patterns of network engagement during task focus for comprehension and arithmetic. Focused comprehension elicits increased activation of semantic regions within the dorsomedial DMN subsystem, plus deactivation of the core DMN, control, and salience networks. In contrast, focused arithmetic is associated with greater engagement of occipital pole and motor cortex, and deactivation of medial occipital, temporal and parietal cortex.

To examine differences in the focus effect between comprehension and arithmetic, we conducted contrast analyses. As shown in Fig. 2B, the left lateral temporal gyrus and IFG showed stronger focus effects during comprehension, whereas bilateral paracingulate gyrus, precuneus cortex, angular gyrus (AG) supramarginal gyrus (SMG), MFG/SFG, frontal pole and right cingulate gyrus showed stronger focus effects during arithmetic.

#### Effects of difficulty in comprehension and arithmetic

Next, we examined whether the neural effects of task focus overlapped with those of difficulty within each domain. In the arithmetic task, regions of motor cortex associated with increased focus also showed greater activation for difficult relative to easy trials, while regions linked to lower focus (characterized by more off-task thought) overlapped with DMN areas that were more active during easy trials (Fig. 2D). This alignment suggests that focus and difficulty in the arithmetic task depend on shared mechanisms: sustaining attention requires engagement of visual–motor systems and suppression of default mode activity, both of which intensify as task demands increase. In contrast, regions associated with increased focus in comprehension overlapped with those more active during easy than hard trials in posterior temporal cortex (Fig. 2C). This implies that sustained focus in reading is not a function of increased effort and might instead reflect successful engagement of temporal lobe semantic regions.

### Study 2: Effects of focus in comprehension versus autobiographical memory recall

Study 1 revealed that sustained focus in externally oriented cognition is associated with different neural mechanisms across semantic and non-semantic tasks: comprehension engaged memory systems while suppressing control regions, whereas higher focus on arithmetic was associated with sensory-motor activity. In Study 2, we asked whether similar effects occur during internally directed cognition, and whether the mechanisms associated with task focus generalize across internal and external modes of thought. Participants alternated between comprehension and autobiographical memory retrieval tasks presented either alone or in competition, allowing us to test whether sustaining focus is associated with a shared neural process of control disengagement, irrespective of the sensory modality or the type of memory organizing cognition.

Twenty-six participants took part in Study 2, which was adapted from a previous task paradigm ^25^. On day 1, participants generated personal memories linked to cue words (e.g., gift) outside the scanner. On day 2, in the scanner, participants were asked to either comprehend the sentences (presented visually or auditorily) or recall their memories for the cue word. The experiment consisted of the following conditions: (1) visual sentences; (2) auditory sentences; (3) retrieve personal memories while ignoring visual sentences; (4) retrieve personal memories while ignoring auditory sentences; (5) retrieve personal memories (no distracting sentence presented; see Fig. 1C for task illustration). After each trial, task focus ratings were collected on a scale of 1 (i.e., not at all focused) to 4 (i.e., highly focused), indexing the extent to which participants were able to focus on each task condition.

#### Behavioral results

Repeated-measures ANOVA on the task focus ratings revealed higher focus in comprehension than recall (*F*(1,25) = 17.06, *p* < .001, *η*^*2*^ = .41). There was a main effect of modality (*F*(1,25) = 7.00, *p* = .014, *η*^*2*^ = .22), and a significant task-by-modality interaction (*F*(1,25) = 7.95, *p* = .009, *η*^*2*^ = .24; Fig. 1D). Post-hoc Bonferroni-corrected *t*-tests revealed higher focus when ignoring visual than auditory sentences during recall (*t*(25) = - 3.36, Bonferroni-corrected *p* = .003), but no modality differences during comprehension (*t*(25) = 0.37).

A comparison of the three recall conditions (i.e., retrieving personal memories while ignoring auditory sentences, ignoring visual sentences, or with no distracting sentences presented), revealed a significant main effect of condition (*F*(2,23) = 20.93, *p* < .001, *η*^*2*^ = .46). Participants were more focused during recall without distraction, relative to ignoring auditory (*t*(25) = 5.36, Bonferroni-corrected *p* < .001) or visual sentences (*t*(25) = 3.94, Bonferroni-corrected *p* = .002).

#### Parametric effect of focus ratings

We performed a univariate fMRI analysis to identify the parametric effects of focus ratings on the BOLD response. In the model, the five conditions (i.e., visual sentence, auditory sentence, autobiographical memory while ignoring visual sentences, autobiographical memory while ignoring auditory sentences, autobiographical memory with no distracting sentence presented) were included as five Explanatory Variables (EV) of interest, along with the parametric effect of focus for each condition (see Materials and Methods). Main effects of each task condition are shown in SI Appendix, Fig. S4.

As shown in Fig. 3, greater focus during auditory comprehension was associated with increased activation in semantic areas, including left anterior superior temporal sulcus/gyrus (aSTS/STG) and posterior middle temporal gyrus (pMTG). Greater focus during visual comprehension was similarly associated with increased activation in STG, as well as in bilateral inferior temporal regions such as ventral occipitotemporal cortex (vOT). Using Neurosynth, we performed meta-analysis to reveal functional associations with these regions. Cognitive decoding of the focus map for auditory comprehension identified terms such as “speech”, “auditory” and “comprehension”, while decoding the map for visual comprehension identified terms such as “reading”, “comprehension”, and “words”. More focus in auditory comprehension was associated with greater activation in the dorsal medial DMN subsystem and auditory network, while stronger focus on visual comprehension elicited greater activation in the dorsal medial DMN subsystem, dorsal attention network (DAN) and auditory network.

**Fig 3.**
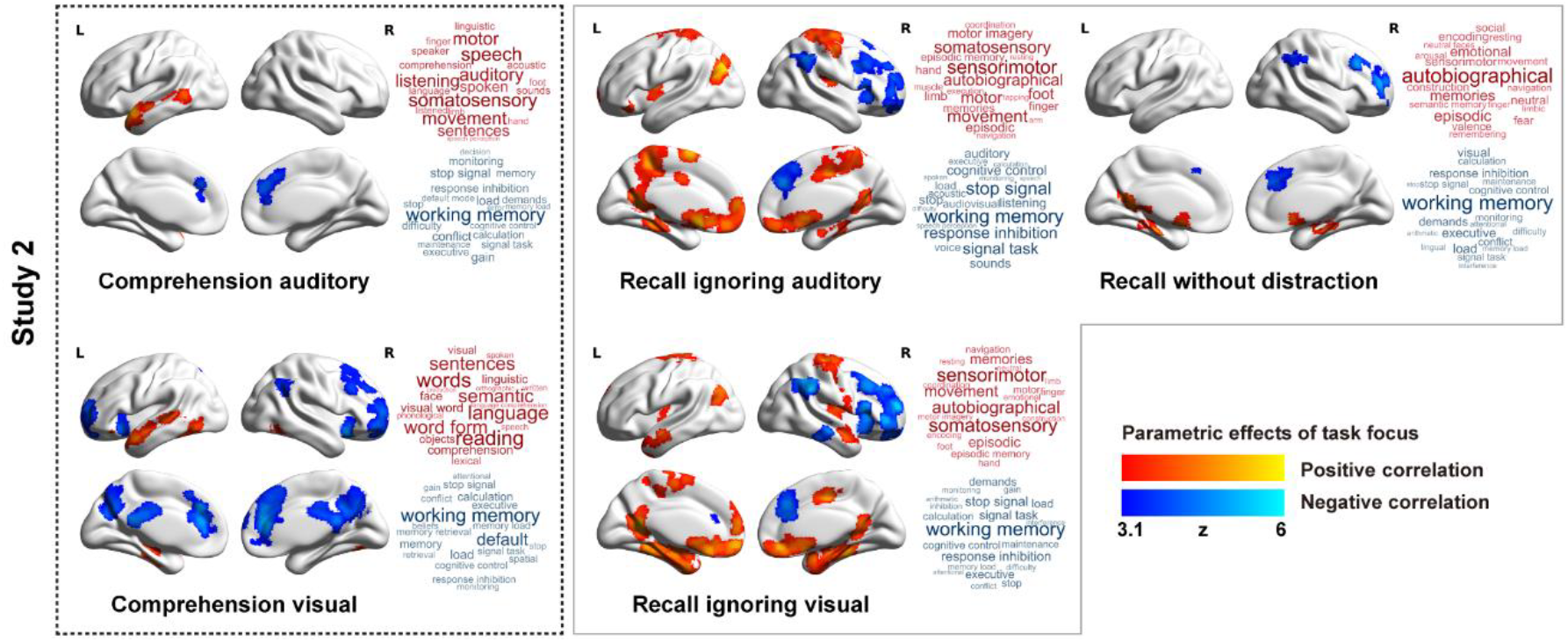
Parametric effects of task focus in Study 2. These brain regions show greater activation (red) or greater deactivation (blue) when participants reported better focus on the task. The word cloud indicates the cognitive decoding of the map using Neurosynth. In the word clouds, the font size of each item illustrates its importance, and the color indicates its association (red = positive correlation, blue = negative correlation). Unthresholded maps of the focus effect, which were used for the Neurosynth decoding analysis, are available at NeuroVault (https://neurovault.org/collections/UXFXQCND/).

Higher focus also elicited greater deactivation in bilateral anterior cingulate and paracingulate cortex for both auditory and visual comprehension. Increased focus on visual comprehension elicited more extensive deactivation in bilateral posterior cingulate, frontal pole, insula, right SFG and MFG. Cognitive decoding of these focus effects for auditory and visual comprehension identified terms such as “working memory” and “inhibition”. Deactivation associated with higher task focus in comprehension therefore largely fell within the control and salience networks, irrespective of input modality, in line with the findings of Study 1.

During memory retrieval, there were positive parametric effects of focus in different brain regions to the comprehension tasks. The effects of focus on memory retrieval were similar for auditory and visual distraction, with increased activation when participants were more focused in bilateral anterior and posterior cingulate, parahippocampal and fusiform cortex, insula, postcentral and precentral gyrus, left superior lateral occipital cortex, SFG and right supplementary motor cortex. Cognitive decoding of these maps identified terms associated with “sensorimotor”, “autobiographical” and “episodic” processing. Higher focus during memory retrieval without external distraction was associated with more constrained activation in bilateral parahippocampal and fusiform cortex, and left precuneus and posterior cingulate; these regions are associated with terms such as “autobiographical”, “episodic” and “memories” when decoded using Neurosynth. This focus-related activation was mainly in the core and medial temporal DMN subsystems and somatomotor network.

Greater focus in memory retrieval was also associated with increased deactivation across all three memory conditions in right AG and SMG, SFG, MFG, IFG, insula and frontal pole. Cognitive decoding revealed that these regions were associated with terms like “working memory”, “response inhibition” and “cognitive control”. This deactivation fell within control and salience networks. When there was no distraction, the overall response was reduced, but activation still fell within the medial temporal DMN subsystem, while the control network and the salience network showed deactivation.

Taken together, increased focus across comprehension and recall elicited greater deactivation in executive networks (i.e., control and salience), with activation patterns that varied according to the process engaged (i.e., comprehension, recall).

### Conjunctions of focus effects across Study 1 and Study 2

Conjunction analyses examined whether the patterns of activation and deactivation during focused comprehension overlapped across datasets. We also asked whether focused comprehension and recall recruited regions that were disengaged by task focus in non-semantic tasks, such as arithmetic. Conjunction analyses used a *z*-statistic threshold of 2.6 and a FWE threshold of *p* < 0.05, in line with previous studies (33, 34). Results are shown in Fig. 4.

**Fig 4.**
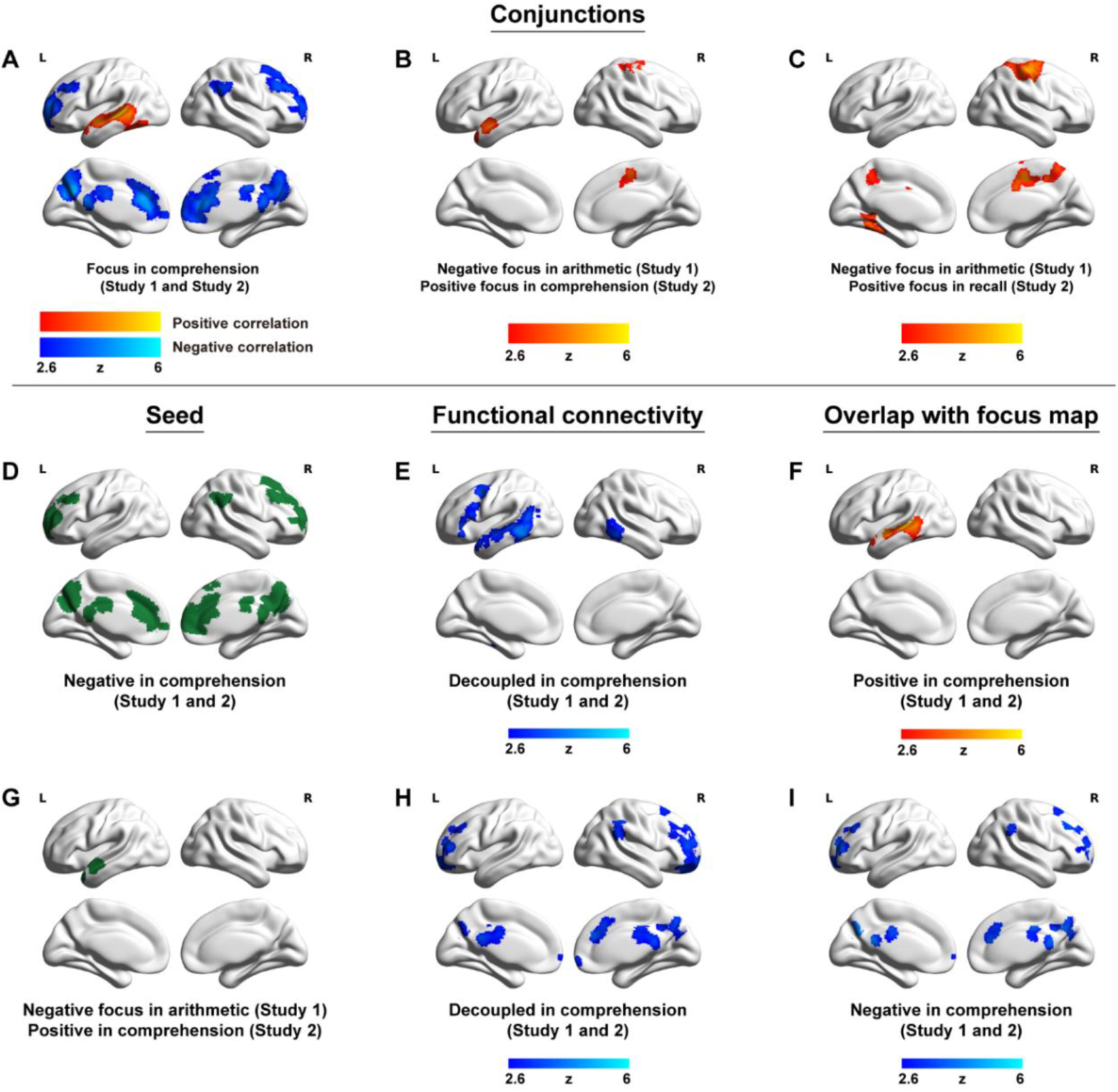
Conjunctions and task based functional connectivity. (A) Formal conjunction of focus effects for comprehension tasks in both Study 1 and Study 2, conducted using FSL “easythresh_conj” tool. Study 1 shows the effect of higher focus in reading, while Study 2 uses the main effect of higher focus in comprehension across visual and auditory modalities. These regions showed greater activation (red) or deactivation (blue) when participants reported better focus during comprehension. (B) The conjunction of negative focus effects during arithmetic in Study 1 and positive focus effects during comprehension in Study 2. Warm colors indicate regions with greater deactivation when participants reported higher focus during arithmetic in Study 1, and greater activation when participants reported higher focus during comprehension in Study 2. (C) The conjunction of negative focus during arithmetic in Study 1 and positive focus effects during recall in Study 2. Warm colors indicate regions with greater deactivation when participants reported higher focus during arithmetic in Study 1, and greater activation when participants reported higher focus during recall in Study 2. (D) Seed included regions showing greater deactivation with higher focus during comprehension across Studies 1 and 2. (E) PPI map from (D, seed), showing regions (blue) with increased decoupling when focus was higher. (F) Overlap of (E) with conjunction focus map, highlighting that these regions (red) also showed greater activation when comprehension focus was high. (G) Seed showing greater activation with higher focus in comprehension (Study 2). (H) PPI map from (G, seed), showing regions (blue) with greater decoupling during high focus. (I) Overlap of (H) with conjunction focus map, highlighting that these regions (blue) also showed greater deactivation when comprehension focus was high. Source data of unthresholded maps in (A) – (C), (E) and (H) are available at NeuroVault (https://neurovault.org/collections/UXFXQCND/).

The conjunction of focused comprehension across Studies 1 and 2 (Fig. 4A) revealed common areas of increased activation in left vOT and MTG/STG, and deactivation in bilateral anterior and posterior cingulate, paracingulate gyrus, precuneus, frontal pole, MFG and right SFG, AG/SMG. When participants were more focused on comprehension, there was a reliable pattern of increased activation in lateral semantic regions, and decreased activation in the core DMN and executive regions.

A conjunction of negative focus during arithmetic in Study 1 and positive focus in comprehension (across modalities in Study 2; Fig. 4B) revealed overlap in left anterior temporal lobe and right precentral/postcentral gyrus and supplementary motor cortex. We compared this conjunction map with a meta-analytic map from Neurosynth (32) for the term ‘semantic’ (1031 contributing studies), which revealed overlap in left ATL (Fig. 4G). Therefore, semantic regions are engaged during focused comprehension, and are suppressed during arithmetic, which does not require access to the semantic store.

The conjunction of negative focus during arithmetic in Study 1 and positive focus during recall in Study 2 (Fig. 4C) further revealed bilateral precuneus, right precentral/postcentral gyrus, SPL, and supplementary motor cortex, and left parahippocampal gyrus/occipitotemporal regions. The left parahippocampal gyrus, occipitotemporal cortex and precuneus overlapped with the Neurosynth meta-analytic map for ‘episodic’ (488 contributing studies). These results indicate that regions associated with episodic memory are engaged during focused recall and suppressed when participants focus on arithmetic.

### Effects of focus on functional connectivity

Using conjunctions that highlighted the distinct roles of control and semantic memory systems in task focus as seeds, we performed psychophysiological interactions (PPI) across the whole brain to investigate variation in connectivity as a function of focus across tasks. We seeded control, salience and default mode network regions that deactivated during more focused comprehension across studies (blue regions in Fig. 4D). We also seeded memory-related regions in ATL (Fig. 4G), which overlapped with the meta-analytic maps for ‘semantic’; activation in this site increased with focus during comprehension and decreased for arithmetic. We constructed separate PPI models for these two seeds and examined the parametric effect of task focus on connectivity in Studies 1 and 2 separately. The PPI models included all the regressors in the basic GLM model, a PPI term for each focus effect, as well as the time series of the seed (for details, See Materials and Methods), using the generalized psychophysiological interaction (gPPI) approach ^45^. The regressors were not orthogonalized. The two datasets revealed similar patterns, and we focus on effects observed reliably across both studies below (see SI Appendix Fig. S5 for all PPI results).

The first seed, showing deactivation in focused comprehension in medial and frontal regions (Fig. 4D), showed greater functional segregation with bilateral MTG/ITG, and left lateral frontal cortex (IFG, precentral gyrus, MFG) when comprehension focus was higher (see Fig. 4E). The overlap (Fig. 4F) of these PPI results with the conjunction map for focus effects in comprehension (brain map in Fig. 4A) demonstrates that control and core DMN areas become increasingly decoupled from regions supporting focused comprehension, suggesting this pattern of disengagement occurs when semantic systems maintain attention with minimal interference.

The second seed, in left ATL, showed greater activation when participants reported higher focus during comprehension (Fig. 4G). As focus increased, the ATL became more functionally segregated from areas in bilateral MFG, frontal pole, precuneus, right anterior cingulate/paracingulate gyrus, SFG, and SMG across Studies 1 and 2 (Fig. 4H). Masking the conjunction focus map with this PPI highlighted the same restricted set of regions largely in control networks (Fig. 4I), which showed deactivation during focused comprehension. These results suggest that focus is associated with functional segregation of language and semantic regions from control and core DMN networks. Together with the first seed, these results demonstrate bidirectional disengagement between semantic and control regions during focused comprehension, regardless of which network is seeded.

## Discussion

Our findings across two independent studies challenge accounts in which sustained focus is primarily a product of executive control. Instead, the neural correlates of task focus were found to vary across cognitive modes, with meaning-based contexts associated with reduced control-network activity and greater functional segregation between control and memory systems. Participants performed externally oriented tasks (comprehension or arithmetic) preceded by autobiographical cues, creating competition between task goals and self-generated thought, and rated their focus after each trial. In Study 1, focus during arithmetic was associated with visual–motor regions, while focused comprehension was associated with greater engagement of semantic regions and reduced activity in control and salience networks. In Study 2, comprehension and autobiographical memory showed convergent patterns: focused cognition coincided with engagement of memory systems and reduced control involvement. In both studies, control and memory systems showed reciprocal separation when participants were more focused.

Functional separation of control and memory systems may be associated with a semantic “flow” state that is relatively independent of executive control ^28, 46^. In this view, coherent thought may be sustained through intrinsic dynamics within memory systems. Rather than requiring continuous external regulation of goal-relevant representations by control networks, these systems may contain structural constraints—such as semantic associations—that allow cognition to proceed along well-established pathways ^47, 48, 49^. Thus, being “on task” during comprehension may reflect a handover from executive control to self-organizing memory systems, with thought sustained by the internal structure of the text’s meaning.

Executive networks are typically engaged during demanding tasks, amplifying relevant information and suppressing distraction ^7, 9, 50, 51, 52^. Executive-failure accounts propose that lapses of attention arise when such control is insufficient, allowing internally generated thought to intrude ^16, 17^. Yet stronger control recruitment does not always predict better performance: more erratic, error-prone (“out-of-the-zone”) behavior can be associated with greater control activation ^53^. Our findings refine this perspective. During arithmetic, higher focus was associated with visual–motor regions, whereas during semantic comprehension and memory retrieval it coincided with reduced engagement of control and salience networks, indicating distinct neural routes to focused cognition.

Behavioral studies indicate that the link between executive control and mind-wandering is more nuanced than simple executive-failure accounts suggest. In non-semantic tasks, mind-wandering typically decreases as difficulty increases, consistent with greater control demands ^23, 54, 55^. In contrast, in semantic tasks such as reading, greater difficulty can increase mind-wandering ^40, 56, 57^, potentially because disruption to semantic prediction and integration promotes internally generated thought. Under such conditions, semantic memory may decouple from sensory input, allowing internally driven associations to dominate over text-related processing. These dissociations suggest that the mechanisms supporting sustained attention differ across cognitive modes, depending on whether cognition is control-driven or memory-driven.

Our neural data help to explain these behavioral patterns and show they cannot be reduced to task difficulty alone. Focused comprehension was associated with reduced engagement of control and salience networks, together with weaker functional coupling between these regions and lateral temporal cortex. If this pattern merely reflected lower executive demands, similar effects would be expected across tasks. Instead, arithmetic showed a partially opposing profile, with higher on-task thought linked to greater engagement of visual–motor systems and stronger deactivation of memory-related regions. Within comprehension, parametric analyses further isolated trial-by-trial variation in subjective focus independent of easy and hard task conditions, indicating that these neural signatures tracked attentional state rather than difficulty per se. Across studies, comprehension and autobiographical retrieval converged in linking higher focus to reduced control involvement despite differing task goals and stimulus contexts. Together, these findings suggest that sustained attention depends on cognitive mode: in some contexts, focus is linked to increased goal-directed processing, whereas in memory-guided cognition it coincides with reduced executive involvement and greater reliance on memory systems. This pattern points to a third attentional mode—neither passive mind-wandering nor effortful control—in which memory representations sustain coherent thought through a flow-like state of control disengagement.

Sustained comprehension across both reading and listening engaged the same heteromodal semantic and language regions in superior and middle temporal gyri. This modality-invariance indicates that task focus depends on representational coherence in long-term memory rather than attentional mechanisms tied to a specific sensory channel. Anterior STG was implicated in sustained focus in comprehension across modalities and in off-task thought during arithmetic, suggesting that this region underpins both task-relevant semantic retrieval and mind-wandering ^27, 58^. This site is proximal to the “semantic hub” proposed to integrate different modalities to give rise to coherent heteromodal concepts ^59^, and it is implicated in both coupled and decoupled aspects of semantic cognition ^60^. The similarity between reading, listening, and off-task thought during arithmetic likely reflects a role for heteromodal semantic representations in all these extended cognitive states, providing a shared basis for maintaining continuity of meaning over time.

We also observed clear dissociations between DMN subsystems across memory states. The dorsomedial DMN, linked to semantic cognition ^61, 62, 63, 64^, was more engaged during focused comprehension across modalities, consistent with its role in conceptual coherence ^47, 48, 49^. In contrast, the medial temporal and core DMN subsystems were more engaged during focused autobiographical memory retrieval: the MT subsystem is implicated in episodic memory and scene construction ^65, 66, 67^, while core DMN activates during autobiographical memory ^25^ and self-relevant cognition ^22, 68^. Despite these differences, control-network suppression overlapped across conditions: dorsomedial prefrontal cortex showed deactivation with higher focus during reading, listening, and autobiographical retrieval, irrespective of distracting stimuli. This convergence suggests that although distinct DMN subsystems support different forms of memory retrieval, focused memory-based cognition consistently involves reduced executive involvement.

Several limitations should be noted. The present studies focused on arithmetic, comprehension, and autobiographical retrieval, and broader sampling of cognitive domains will be needed to determine how widely these modes generalize. Because fMRI provides correlational measures of neural activity, the observed patterns of network engagement and decoupling do not by themselves establish causal mechanisms ^69, 70^. In addition, self-reported focus likely reflects multiple processes, including motivation, interest, effort, and subjective absorption ^19, 39, 71, 72^. Future work combining causal perturbation methods and temporally resolved imaging could clarify how control and memory systems interact across tasks.

The ability to maintain coherent, goal-directed thought without continuous executive supervision may be highly adaptive. Executive control is cognitively demanding and likely metabolically costly ^73, 74^; by leveraging the self-organizing properties of long-term memory, the brain may sustain focus on meaningful information such as language or autobiographical memories more efficiently. This perspective suggests that when past experience supports cognition, focus may often involve fluent processing within memory systems, with top-down control serving primarily as intermittent guidance or error correction rather than a continuous driver of thought.

## Methods

### Participants

In Study 1, we recruited 35 participants (26 females, mean age = 20 years, range = 18-29 years). All were right-handed native English speakers, and had normal or corrected vision. One was excluded because of big head movement (relative head motion > 0.3 mm). Therefore, the final sample of this analysis included 34 participants (25 females, mean age = 20 years, range = 18-29 years).

In Study 2, we recruited 30 participants (24 females, mean age = 20 years, range = 18-23 years). All were right-handed native English speakers, and had normal or corrected vision, except for one who had amblyopia in the right eye. Four participants were excluded from the analysis, because there was insufficient data to model the parametric effect of task focus after excluding runs in which all trials had the same demeaned task focus rating for at least one of the conditions. Therefore, the final sample included 26 participants (20 females, mean age = 20 years, range = 18-23 years).

In both studies, none of the participants had any history of neurological impairment, diagnosis of learning difficulty or psychiatric illness. All provided written informed consent prior to taking part. Participants were recruited through participant pools and paid for their time or awarded course credits. Ethical approval was obtained from the Research Ethics Committees of the Department of Psychology and York Neuroimaging Centre, University of York (Project number: Study 1 - P1501; Study 2 - P1406).

### Task Design and procedure

Both Studies 1 and 2 were adapted from ref. ^25^ . In Study 1, we employed a within-subjects 2 × 2 design manipulating task (Comprehension vs. Arithmetic) and difficulty (Easy vs. Hard). Participants generated thoughts linked to cue phrases (e.g., “Lottery win”) outside the scanner. Inside the scanner, participants retrieved personal thoughts from cue phrases for 3 s, and then either read factual sentences or added up numbers for 9 s (see Fig. 1A for task illustration). After each comprehension or arithmetic trial, participants provided ratings on a scale from 1 (i.e., thinking about other things) to 7 (i.e., focusing on the meaning of the sentence or the ongoing addition) within 2 s, which measured the extent to which participants were focused on the current task.

Study 2 took place across two days. On day 1, participants generated personal memories linked to cue words (e.g., gift) outside the scanner. On day 2, inside the scanner, participants brought to mind their detailed personal memories related to a written and spoken autobiographical memory cue word for 2 s. Following the task instruction (i.e., COMPREHEND or RECALL) presented for 1 s, participants were asked to either comprehend the sentences (presented visually or auditorily), or recall their memories for the cue word for an average of 9 s. The experiment consisted of the following conditions: (a) visual sentences; (b) auditory sentences; (c) retrieve personal memories while ignoring visual sentences; (d) retrieve personal memories while ignoring auditory sentences; (e) retrieve personal memories (no distracting sentence presented; see Fig. 1C for task illustration). After each trial, task focus ratings on a scale of 1 (i.e., not at all focused) to 4 (i.e., highly focused) were collected within 2 s to index the extent to which participants were able to focus on each task condition. For more information, refer to SI Appendix, Supplemental Methods.

### Materials

Study 1 included 50 paired topic sentences, which therefore resulted in two experiment versions, each containing 25 easy and 25 hard sentences. The versions were counterbalanced across participants. These sentences contained accessible factual content on non-emotive topics, with easy texts selected from BBC Bitesize and hard texts selected from Wikipedia. For example, the topic MYTH included an easy version, “Myths are ancient tales filled with magical creatures, gods and mystery that are passed down the generations and not based on facts or reality”, whereas the hard version was, “Myth is a genre of folklore or theology consisting primarily of narratives that play a fundamental role in society such as foundational tales”. We balanced the number of characters across the easy and hard conditions (for all paired *t*-test, Bonferroni-corrected *p* > .05), to ensure that the visual loadings were comparable across conditions. Overall, the easy sentences were rated as significantly easier to understand, more interesting, more frequent, and they contained more words than the hard sentences (for detailed information, refer to SI Appendix, Supplemental Methods). For the arithmetic task, the easy condition consisted of additions of zeros, whereas the hard condition involved additions with random numbers between 1 and 4. To induce a mind wandering state prior to each comprehension or arithmetic trial, we presented a cue phrase before each sentence. A total of 25 cue phrases were selected from the positive set used in ref. ^75^ (e.g., “Stars at night”, “Lottery win”). The same cue phrases were used across all conditions and were counterbalanced across runs to ensure that no phrase was repeated within the same run.

Study 2 presented participants with 96 sentences, all of which were preceded by a cue word. The sentences were selected using 96 highly frequent and concrete nouns as key words as a search term in Wikipedia to identify text that described largely unfamiliar facts about each item (sentence length: Mean ± SD = 20.33 ± 1.54 words). These sentences contained accessible factual content, on non-emotive topics; for example, “Posters are used for reproductions of artwork, particularly famous works, and are cheaper compared to the original” for the keyword POSTER. Overall, the sentences were unfamiliar, easy to understand and did not differ in familiarity and comprehensibility between conditions (for detailed information, refer to SI Appendix, Supplemental Methods). The cue words were 60 highly frequent and concrete nouns and were repeated twice across the whole experiment. The cue words were matched for lexical frequency and concreteness across conditions. The full set of sentences for both studies 1 and 2 is available on OSF (https://osf.io/cqbhv/overview?view_only=ea533ec03ad3471a8336cc32107c15f6).

### fMRI data acquisition

Whole brain structural and functional MRI data were acquired using a 3T Siemens MRI scanner utilizing a 64-channel head coil, tuned to 123 MHz at the York Neuroimaging Centre, University of York. We used a multiband-multi-echo (MBME) EPI sequence with the following parameters: TR=1.5 s; TEs = 12, 24.83, 37.66 ms; 48 interleaved slices per volume with slice thickness of 3 mm (no slice gap); FoV = 24 cm (resolution matrix = 3x3x3; 80x80); 75° flip angle; volumes per run below; 7/8 partial Fourier encoding and GRAPPA (acceleration factor = 3, 36 ref. lines; multi-band acceleration factor = 2). Structural T1-weighted images were acquired using an MPRAGE sequence (TR = 2.3 s, TE = 2.26 s; voxel size = 1x1x1 isotropic; 176 slices; flip angle = 8°; FoV= 256 mm; interleaved slice ordering). In Study 1, tasks were presented across four runs, each lasting 8.5 min, with 340 volumes in each run. In Study 2, tasks were presented across four runs, each lasting 9.35 min, with 374 volumes in each run.

### Multi-echo data pre-processing

The same preprocessing pipeline was applied to both Study 1 and Study 2. A MBME sequence was used to optimize signal from the anterior and medial temporal regions, while maintaining optimal signal across the whole brain ^76^. We used TE Dependent ANAlysis (TEDANA; version v23.0.1) to combine the images ^77, 78, 79^. Anatomical pre-processing (fsl_anat; https://fsl.fmrib.ox.ac.uk/fsl/fslwiki/fsl_anat) included re-orientation to standard MNI space (fslreorient2std), automatic cropping (robustfov), bias-field correction (RF/B1 – inhomogeneity-correction, using FAST), linear and non-linear registration to standard-space (using FLIRT and FNIRT), brain extraction (using FNIRT, BET), tissue-type segmentation (using FAST) and subcortical structure segmentation (FAST). The multi-echo data were preprocessed using AFNI (https://afni.nimh.nih.gov/), including de-spiking (3dDespike), slice timing correction (3dTshift; heptic interpolation), and motion correction (3dvolreg applied to echo 1 to realign all images to the first volume; these transformation parameters were then applied to echoes 2 and 3; cubic interpolation).The script used to implement the preprocessing TEDANA pipeline is available at OSF (https://osf.io/cqbhv/overview?view_only=ea533ec03ad3471a8336cc32107c15f6).

### fMRI data analysis

Individual level analysis was conducted using FSL-FEAT version 6 (FMRIB’s Software Library, www.fmrib.ox.ac.uk/fsl; ref. ^80, 81, 82^). Denoised optimally-combined time series data were submitted to FSL. Pre-processing included high-pass temporal filtering (Gaussian-weighted least-squares straight line fitting, with sigma = 100s), linear co-registration to the corresponding T1-weighted image and to MNI152 standard space ^83^, spatial smoothing using a Gaussian kernel with full-width-half-maximum (FWHM) of 6 mm, and grand-mean intensity normalization of the entire 4D dataset by a single multiplicative factor. Pre-processed time series data were modelled using a general linear model correcting for local autocorrelation ^84^.

In Study 1, six Explanatory Variables (EV) of interest and four of no interest were modelled using a double-Gaussian hemodynamic response gamma function. The six EVs of interest were as follows: (1) easy comprehension; (2) hard comprehension; (3) easy arithmetic; (4) hard arithmetic; (5-6) task focus effect of comprehension and arithmetic as a parametric regressor. EVs of no interest were: (7) memory cue phrase; (8) fixation before sentences or numbers; (9) fixation before the rating question; (10) rating decision period. In Study 2, ten Explanatory Variables (EV) of interest and four of no interest were modelled using a double-Gaussian hemodynamic response gamma function. The ten EVs of interest were as follows:

(1) visual sentence; (2) auditory sentence; (3) autobiographical memory while ignoring visual sentences; (4) autobiographical memory while ignoring auditory sentences; (5) autobiographical memory with no distracting sentence presented; (6-10) task focus effects for each of the five experimental conditions as a parametric regressor. EVs of no interest were: (11) memory cue word; (12) task instruction; (13) fixation (the period before the rating question); (14) rating decision period. No motion parameters were included in the models, as the data had already been denoised as part of the TEDANA pipeline ^79^. EVs for each condition commenced at the onset of the first word of the sentence or the first number, with EV duration set as the presentation time (e.g., 9 s in Study 1, or the entire length of the sentence in Study 2). The parametric EVs for the effect of task focus had the same onset time and duration for each trial in the EVs corresponding to the comprehension and arithmetic trials, and the demeaned task focus rating as the weight. Runs in which all trials within a condition had the same task focus rating value were excluded from analyses, since it was impossible to model the effects of task focus. This resulted in the removal of four participants’ data in Study 2; on average, more than 77% of the data from Study 2 was retained across participants. The fixation period between trials provided the implicit baseline. In the group-level analysis, clusters were corrected using a threshold of *z* = 3.1 to define contiguous clusters and a familywise-error-corrected (FWE) significance level of *p* < .05.

Psychophysiological interaction (PPI) analyses were conducted separately for Study 1 and Study 2 using seeds derived from conjunction analyses. The model included all the regressors from the basic GLM model described above, a PPI term for each experimental focus effects, as well as the time series of the seed, using the generalized psychophysiological interaction (gPPI) approach ^45^. The regressors were not orthogonalized. All Brain figures were visualized using BrainNet Viewer ^85^.

## Acknowledgments

We thank all the staff in the York Neuroimage Center for their supporting in fMRI scanning. This project was supported by the European Research Council (Project ID: 771863 – FLEXSEM to E.J.), a Horizon Europe Guarantee MSCA Postdoctoral Fellowship by UK Research and Innovation (EP/Y014367/1 to L.M.-M.) and a China Scholarship Council (CSC) Scholarship (No. 202206870011 to C.C.).

## Author Contributions

C.C., K.K.-R., M.Z., J.S., and E.J. designed research; C.C., K.K.-R., and L.M.-M. performed research; C.C., K.K.-R., and X.S. analyzed data; C.C., K.K.-R., and E.J. wrote the paper.

## Data Availability Statement

We have sufficient consent for public sharing of the data. The experiment materials, scripts for task, and behavioural data are made available on OSF: https://osf.io/cqbhv/overview?view_only=ea533ec03ad3471a8336cc32107c15f6. The grouplevel fMRI data, including unthresholded maps, are on Neurovault: https://neurovault.org/collections/UXFXQCND/. Given the very large files sizes involved in the fMRI dataset, these data are not on a public repository but will be made available to researchers following an email request to the York Neuroimaging Centre’s Research Ethics Committee, and details on how to access the data are given in the manuscript.

## Ethics Statement

All participants provided informed consent prior to participation, and all procedures were conducted in accordance with the approved ethical guidelines. Ethical approval was obtained from the Research Ethics Committees of the Department of Psychology and York Neuroimaging Centre, University of York (Project number: Study 1 - P1501; Study 2 - P1406).

