## Supplementary material for "Distinct neural architectures prevent off-task thought in different task contexts": SI Appendix

**Supporting Information Text**

**Supplemental Methods**

**Procedure.**

**Study 1:** Participants first generated their personal memories linked to autobiographical cue phrases outside the scanner. They were asked to think about a personal memory involving someone they knew, a show, song, or event, future plans or personal ideas. They then completed twelve practice trials to familiarize themselves with the paradigm. In the scanner, as shown in Fig. 1A, each trial started with a fixation cross (1-3 s) at the center of the screen, followed by an autobiographical cue phrase lasting 3 s. During the presentation of the cue phrase, participants brought to mind their detailed personal memories related to this item. Next, a jittered fixation (1-3 s) was presented before the sentences or numbers.

Following the fixation, the main trial began with presentation of stimuli lasting for 9 s. Stimuli consisted of either sentences or numbers. Sentences were presented on 15 successive slides, with each one lasting 600 ms, combining short words on a single slide (e.g., "such as") or presenting articles and conjunctions together with nouns (e.g., "a minute"). Numbers were presented on every other slide, with a blank screen in between, resulting in a total of eight number slides. Each trial was followed by a jittered fixation (1-3) and a task focus rating question. Participants were required to rate their task focus on a 7-point scale of from 1 (i.e., think about other things) to 7 (i.e., think about the meaning of the sentence or the ongoing addition of the numbers) within 2 s. Responses were made using two button boxes, with the left hand indicating lower ratings and the right hand indicating higher ratings.

Each condition contained 25 trials and the stimuli were presented across four runs, with each containing 25 trials. Each trial lasted on an average of 20 s. The whole experiment lasted 32 minutes. Trials were presented using a mini-block design: within each run, all trials of the same condition were presented consecutively in a pseudorandom order, and the order of conditions was counterbalanced across runs.

**Study 2:** This study occurred across two consecutive days. On day 1, participants generated their personal memories linked to autobiographical cue words (e.g., sea) outside the scanner. They were asked to identify specific events that they were personally involved in and to provide as much detail about these events as they could, including when and where the event took place, who was involved, what happened, and the duration. They were asked to type these details into a spreadsheet, which ensured that comparable information was recorded for different cue words.

On day 2, before entering the scanner, participants were asked to review their generated memories to ensure they were easily accessible when presented with the cue word in the scanner. Next, they performed ten practice trials to familiarize themselves with the paradigm. In the scanner, participants engaged in internal recall of their memories when presented with a cue word (i.e., participants did not speak in the scanner), and also read and listened to factual sentences in a comprehension task. As shown in Fig. 1C, each trial started with a fixation cross (1-3 s) in the center of the screen, followed by a written and spoken autobiographical memory cue word lasting 2 s. During the presentation of the cue word, participants brought to mind their detailed personal memories relating to this item. Next, the task instruction (i.e., COMPREHEND or RECALL) was presented for 1 s, in either a visual (i.e., text) or auditory (i.e., voice) format. Following the task instruction, the main trial began, with presentation of stimuli lasting for 7.8-10.2 s, with an average of 9 s. Stimuli consisted of either (1a) visual or (1b) auditory sentences or (2a) a series of x's (visual) or (2b) white noise (auditory). The visual sentences were presented on 13-17 successive slides, with each one lasting 600 ms, combining short words on a single slide (e.g., "such as") or presenting articles and conjunctions together with nouns (e.g., "the heat"). The auditory sentences were presented naturalistically, with the sentence unfolding as the speaker intended. The visual instructions and sentences were presented together with white noise, and the auditory instructions and sentences were presented together with letter strings (e.g., XXXXX), which varied in length (i.e., XXX ; XXXXXX, etc), so that sensory inputs were presented in both visual and auditory channels across all conditions. On memory recall trials without a conflicting sentence, the task instructions were presented visually on half of the trials and auditorily on the other half, followed by both letter strings and white noise. On memory recall trials, participants were instructed to keep thinking about their autobiographical memory, in as much detail as possible, until the end of the trial. Each trial was followed by a jittered fixation (1-3) and a task focus rating question. Participants were required to rate their task focus on a scale of 1 (i.e., not at all focused) to 4 (i.e., highly focused) within 2 s, using a button press to respond.

Each condition contained 24 trials; stimuli were presented across four runs, with each containing 30 trials. Each trial lasted 14.8-21.2 s with an average of 18.0 s. The whole experiment lasted 36 minutes. Trials were presented in a pseudorandom order, ensuring that trials from the same experimental condition were not presented consecutively more than twice, and the memory cue word of each trial was only presented once in each run (the memory cue word was presented a maximum of two times across the whole experiment).

**Materials.**

**Study 1:** This study included 50 paired topic sentences, which therefore resulted in two experiment versions, each containing 25 easy and 25 hard sentences. The versions were counterbalanced across participants. These sentences contained accessible factual content on non-emotive topics, with easy texts selected from BBC Bitesize and hard texts selected from Wikipedia. For example, the topic MYTH included an easy version, “Myths are ancient tales filled with magical creatures, gods and mystery that are passed down the generations and not based on facts or reality”, whereas the hard version was, “Myth is a genre of folklore or theology consisting primarily of narratives that play a fundamental role in society such as foundational tales”. The full set of stimuli used is available on OSF. We collected ratings for the sentences used in this study from separate cohorts of participants (i.e., individuals who did not take part in the fMRI experiment). Participants rated the difficulty (n=24), and interestingness (n=22) of the sentences on a 7-point scale, ranging from 1 (i.e., “extremely easy”; “not at all interesting”) to 7 (i.e., “extremely difficult”; “extremely interesting”). Overall, the easy sentences were rated as significantly easier to understand, more interesting, more frequent (SUBTLEX_US word frequency database; ref. [1]), and they contained more words than the hard sentences (Bonferroni-corrected $p < .05$). There was no difference in concreteness (ref. [2]; all $F < .10$ for cue words). We balanced the number of characters across the easy and hard conditions (all paired t-test, Bonferroni-corrected $p > .05$), to ensure that the visual loadings were comparable across conditions.

For the arithmetic task, the easy condition consisted of additions of zeros, whereas the hard condition involved additions with random numbers between 1 and 4.

To induce a mind wandering state prior to each comprehension or arithmetic trial, we presented a cue phrase before each sentence. A total of 25 cue phrases were selected from the positive set used in ref. [3] (e.g., “Stars at night”, “Lottery win”). The same cue phrases were used across all conditions and were counterbalanced across runs to ensure that no phrase was repeated within the same run.

**Study 2:** This study presented participants with 96 sentences, all of which were preceded by a cue word. The sentences were selected using 96 highly frequent and concrete nouns as key words as a search term in Wikipedia to identify text that described largely unfamiliar facts about each item (sentence length: Mean ± SD = 20.33 ± 1.54 words). These sentences contained accessible factual content, on non-emotive topics; for example, “Posters are used for reproductions of artwork, particularly famous works, and are cheaper compared to the original” for the keyword POSTER. The full set of sentences is available on OSF (https://osf.io/cqbhv/overview?view_only=ea533ec03ad3471a8336cc32107c15f6). The sentences were divided into four sets and assigned to the following conditions: (1) read sentences; (2) listen to sentences; (3) retrieve personal memories while ignoring visual sentences; (4) retrieve personal memories while ignoring auditory sentences. We collected ratings for the sentences used in this study from separate cohorts of participants (i.e., they did not take part in the fMRI experiment). Participants rated the familiarity (n=16), and comprehensibility (n=20) of the sentences, on a 7-point scale, from 1 (i.e., “completely unfamiliar”; “very hard to understand”) to 7 (i.e., “extremely familiar”; “very easy to understand”). Overall, the sentences were unfamiliar (familiarity: Mean ± SD = 4.23 ± 1.15), easy to understand (comprehensibility: Mean ± SD = 6.12 ± 0.43) and did not differ in familiarity and comprehensibility between conditions (all $F < .38$).

Each sentence was preceded by a cue word, to induce autobiographical memory recall. These cue words were 60 highly frequent and concrete nouns. The autobiographical memory cues were repeated twice across the whole experiment. Spoken autobiographical memory cues and sentences were recorded by a male native British English speaker, using Audacity (http://audacity.sourceforge.net/), in a sound-attenuated room. The stimuli were normalized for volume and power in Audacity, producing a level of −3 dB FS. The MRI auditory stimulus system (MR Confon mkII+, www.mr-confon.de/en/products.html) was used to give a maximum presentation level of 80–90 dB SPL.

The key words and cue words were matched for key psycholinguistic properties across conditions: they did not differ in lexical frequency (SUBTLEX_US word frequency database; ref. [1]) or concreteness (ref. [2]; all $F < .10$ for cue words). In addition, there were no significant differences between the four sentence conditions in lexical frequency or concreteness for all the words contained in the sentences (all $F < 0.44$).

**Analysis.**

Parametric effects of task focus were decoded using the online tool Neurosynth [4]. Neurosynth (RRID: SCR_006798) is an automated meta-analysis tool that uses text-mining approaches to extract terms from neuroimaging articles that typically co-occur with specific peak coordinates of activation. It can be used to generate a set of terms frequently associated with a spatial map. Word clouds were generated using Python. We excluded terms referring to neuroanatomy (e.g., “inferior” or “sulcus”), and repeated terms (e.g., “semantic” and “semantics”). The size of the word in the word cloud reflects the frequency of that term across other studies, and the color indicates its relationship with focus rating in our study (i.e., red = positive correlation with task focus, blue = negative correlation with task focus).

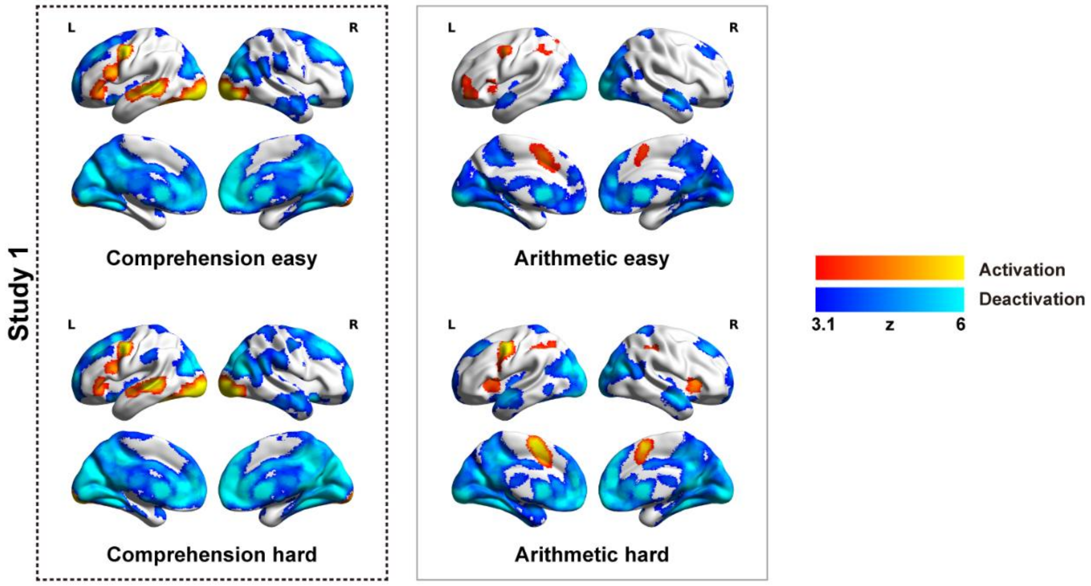


**Fig. S1.** Univariate activation results for comprehension and arithmetic in study 1 (N = 34). Warm colors = task > implicit baseline. Cold colors = task < implicit baseline. All maps were cluster corrected at $z > 3.1$, $p < .05$.

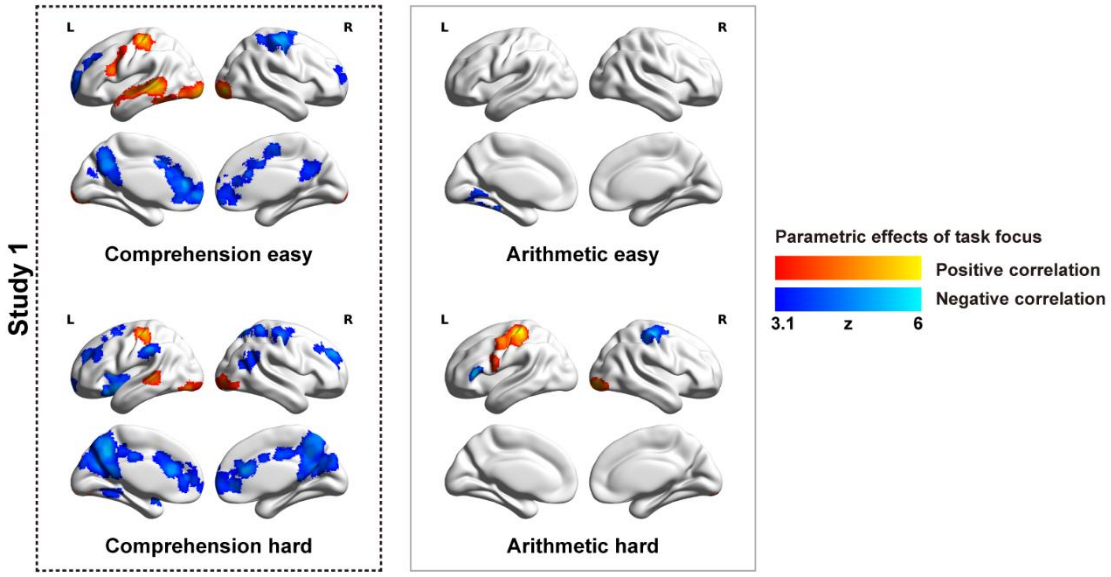


**Fig. S2.** Parametric effects of task focus for easy and hard comprehension and arithmetic in study 1. In this analysis, eight Explanatory Variables (EV) of interest and four of no interest were modelled using a double-Gaussian hemodynamic response gamma function: 1. easy comprehension; 2. hard comprehension; 3. easy arithmetic; 4. hard arithmetic; 5-8. parametric effect of focus for easy and hard comprehension and arithmetic; 9. memory cue phrase; 10. fixation before sentences or numbers; 11. fixation before the rating question; 12. rating decision period. 35 participants were recruited; one was excluded because of big head movement, and another was excluded due to a lack of variation in focus ratings during easy arithmetic task across all runs. Therefore, 33 participants were included in this group-level analysis. These brain regions show greater activation (red) or greater deactivation (blue) when participants reported higher task focus. Results were cluster corrected at $z > 3.1$, $p < .05$.

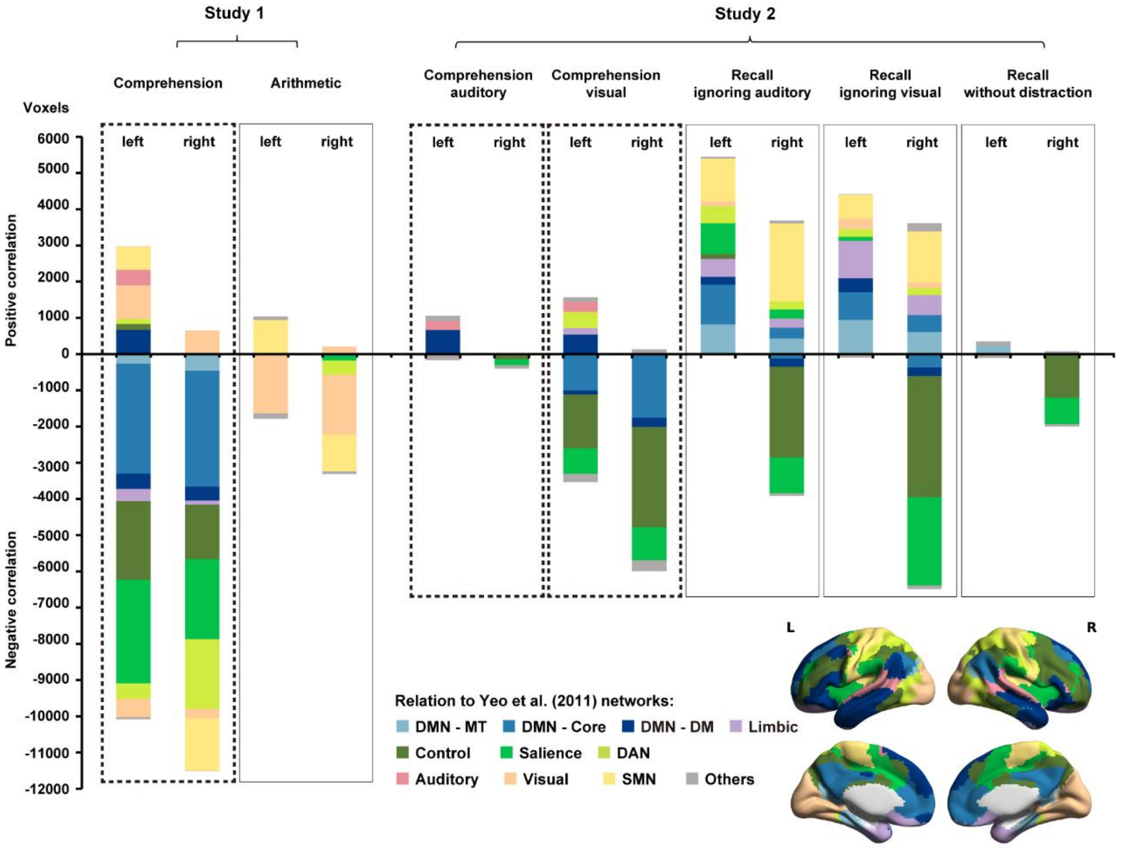


**Fig. S3.** Network analysis of focus effects in Study 1 and 2. The chart shows counts of voxels per network that were significantly correlated with task focus ratings. Top: networks showing greater activation with higher task focus. Bottom: networks showing greater deactivation with higher task focus (see Fig. 2 and Fig. 4). Networks with fewer than 100 activated/deactivated voxels were collapsed into the "Others" category. DMN, default mode network; MT, medial temporal; DM, dorsal medial; DAN, dorsal attention network; SMN, somatomotor network.

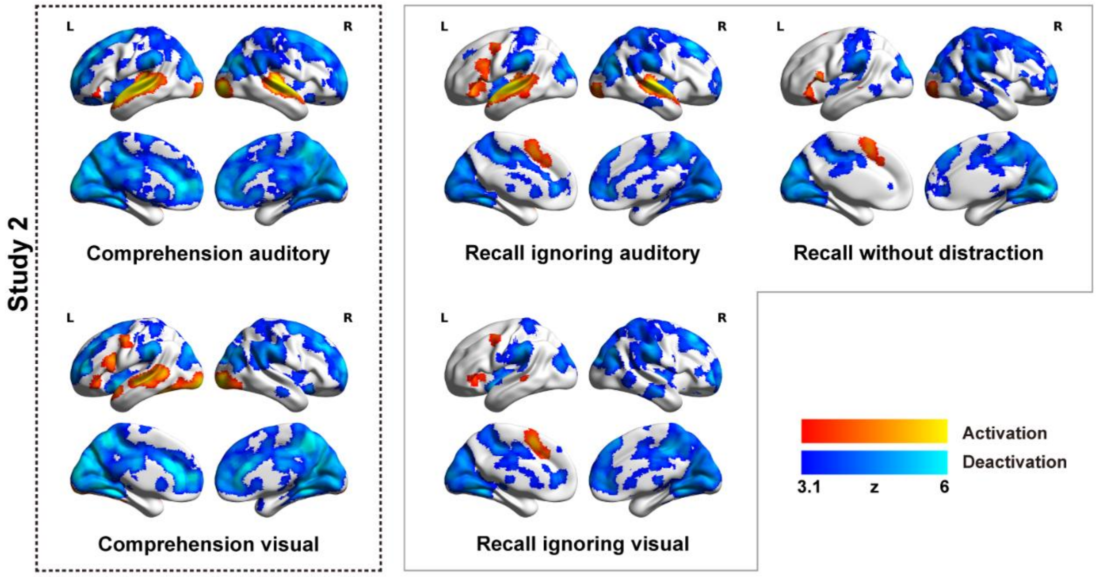


**Fig. S4.** Univariate activation results for comprehension and autobiographical memory retrieval in study 2 (N = 26). Warm colors = task > implicit baseline. Cold colors = task < implicit baseline. All maps were cluster corrected at $z > 3.1$, $p < .05$.

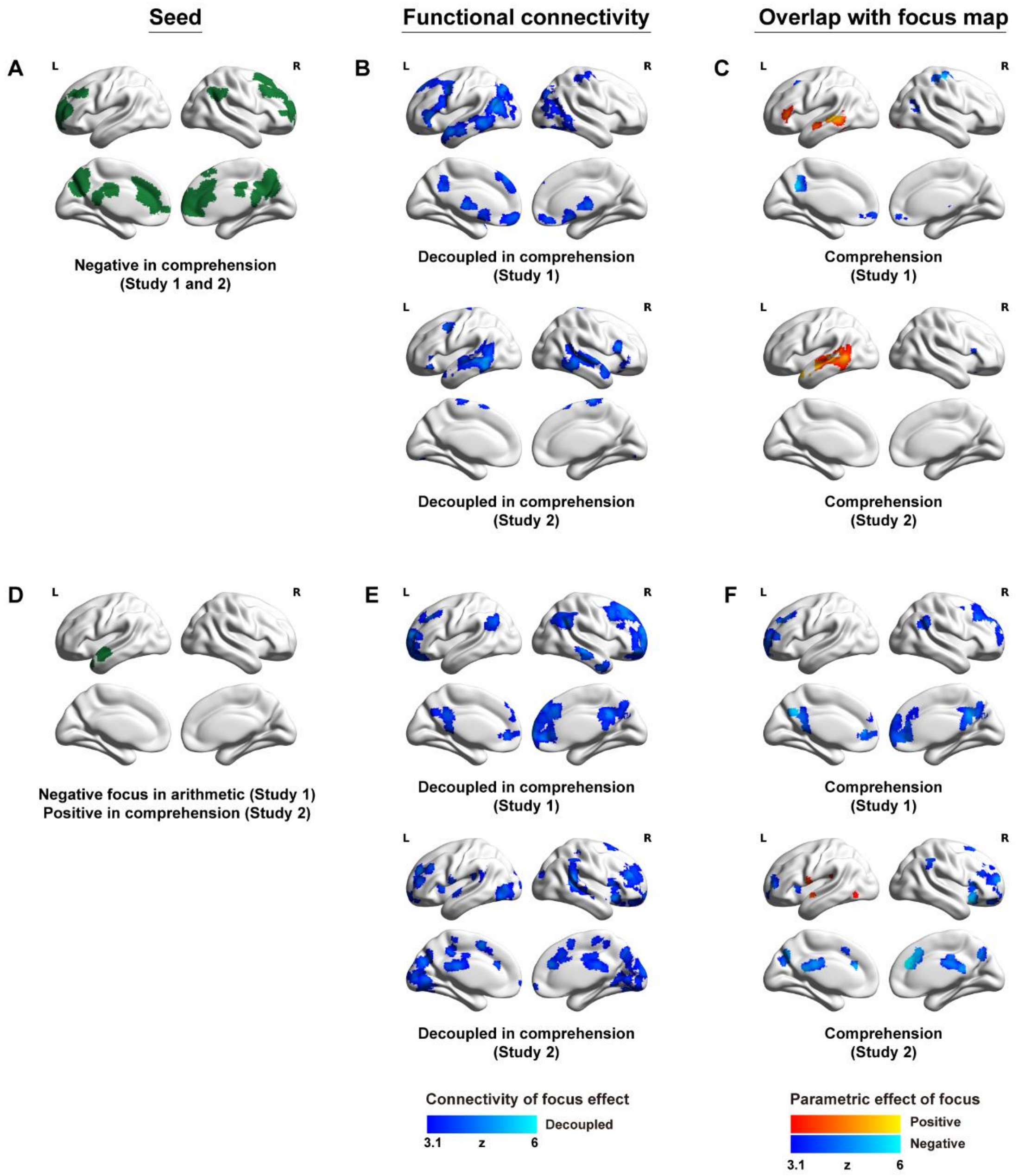


**Fig. S5.** Task-based functional connectivity. (A) Seed showing greater deactivation when participants reported higher focus during comprehension across studies 1 and 2. (B) The functional connectivity maps seeded from (A). These regions (blue) showed greater decoupling from the seed when higher focus was reported during comprehension in study 1 or study 2. (C) The overlap regions obtained by masking focus maps by regions in (B), showing greater activation (red) or deactivation (blue) when greater focus was reported during comprehension. (D) Seed showing greater activation when participants reported higher focus during comprehension in study 1. (E) The functional connectivity maps seeded from (D). These regions (blue) showed greater decoupling from the seed when higher focus was reported during comprehension. (F) The overlap regions obtained by masking the focus maps by regions in (E), showing greater deactivation (blue) when greater focus was reported during comprehension. Results were cluster corrected at $z > 3.1$, $p < .05$.

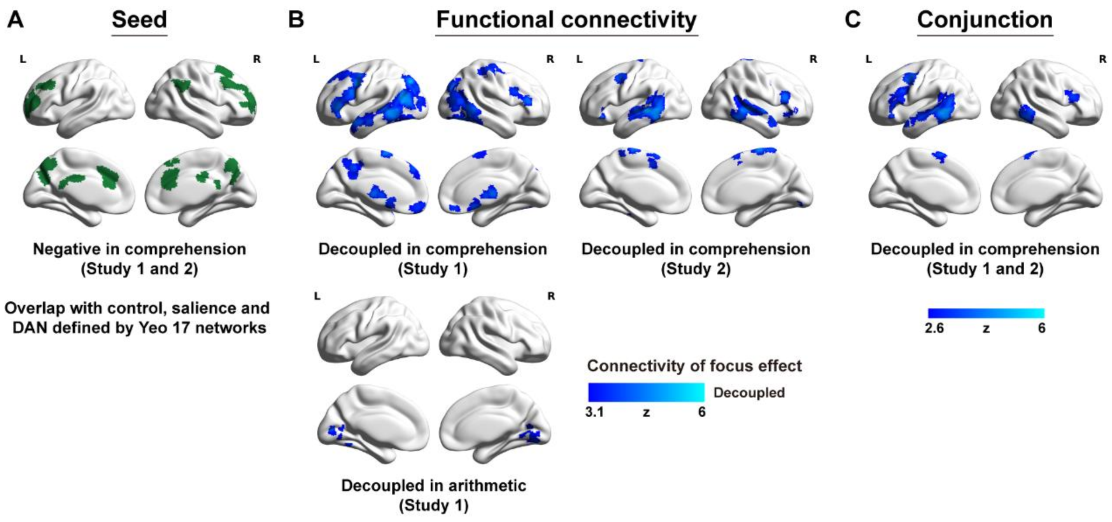


**Fig. S6.** Task-based functional connectivity seeding from Control/Salience/DAN regions (A). (A) Seed region showing greater deactivation when participants reported higher focus during comprehension across studies 1 and 2. This seed was generated from the conjunction of comprehension across studies 1 and 2, masked by the control, salience, and dorsal attention (DAN) networks defined by ref. [5]. (B) Functional connectivity maps seeded from (A). These regions (blue) showed greater decoupling from the seed when higher task focus was reported. Results were cluster corrected at $z > 3.1$, $p < .05$. (C) The conjunction of PPI results during comprehension across studies 1 and 2. These regions (blue) showed greater decoupling from the seed when higher focus was reported during comprehension. The conjunction result was cluster corrected at $z > 2.6$, $p < .05$.

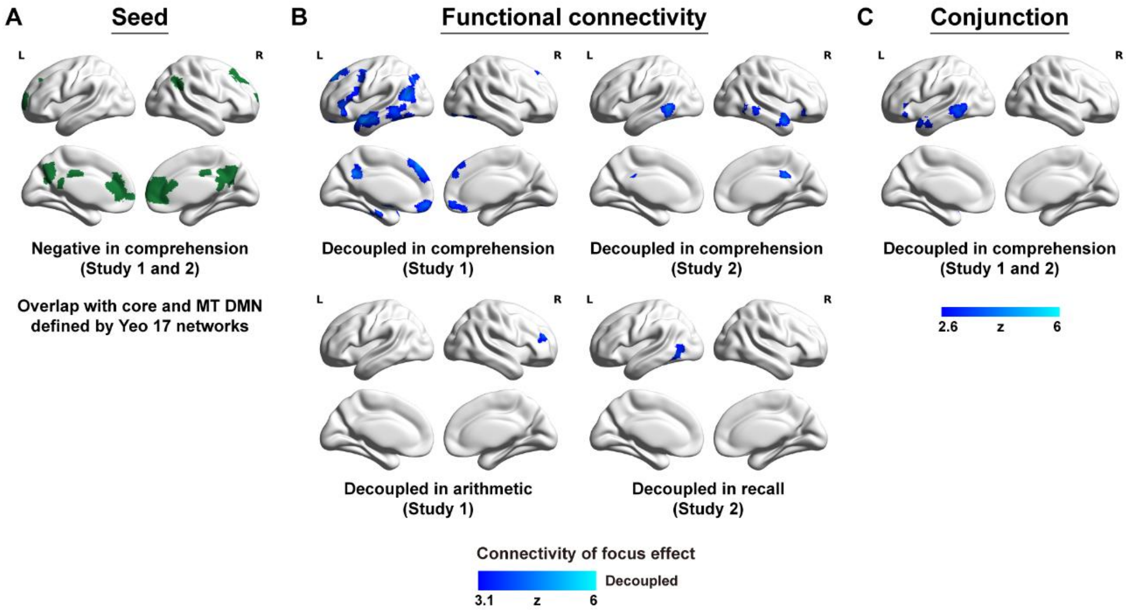


**Fig. S7.** Task-based functional connectivity seeding from Core/MT DMN regions (A). (A) Seed region showing greater deactivation when participants reported higher focus during comprehension across studies 1 and 2. This seed was generated from the conjunction of comprehension across studies 1 and 2, masked by the core and medial temporal (MT) DMN defined by ref. [5]. (B) Functional connectivity maps seeded from (A). These regions (blue) showed greater decoupling from the seed when higher task focus was reported. Results were cluster corrected at $z > 3.1$, $p < .05$. (C) The conjunction of PPI results during comprehension across studies 1 and 2. These regions (blue) showed greater decoupling from the seed when higher focus was reported during comprehension. The conjunction result was cluster corrected at $z > 2.6$, $p < .05$.

**Table S1.** Study 1 linguistic properties of sentences, mean (SD).

| Sentences | Word count | Character count | Difficulty | Frequency | Concreteness | Interestingness |
|---|---|---|---|---|---|---|
| Version 1 - comprehension - easy | 21 (3) | 100 (13) | 1.83 (.37) | 4.15 (.28) | 2.76 (.23) | 3.25 (.91) |
| Version 1 - comprehension - hard | 19 (2) | 103 (12) | 4.22 (.68) | 3.67 (.41) | 2.62 (.20) | 2.5 (.74) |
| Version 2 - comprehension - easy | 22 (2) | 101 (9) | 1.85 (.36) | 4.19 (.37) | 2.73 (.18) | 3.34 (.79) |
| Version 2 - comprehension - hard | 19 (2) | 104 (14) | 4.43 (.76) | 3.71 (.46) | 2.58 (.28) | 2.41 (.67) |

**Table S2.** Study 2 linguistic properties of cue words and sentences, mean (SD).

| Stimuli set | Frequency | Concreteness | Familiarity | Comprehensibility |
|---|---|---|---|---|
| (i) autobiographical memory cues | 2.83 (.50) | 4.73 (.29) | / | / |
| (ii) autobiographical memory cues | 2.86 (.50) | 4.69 (.43) | / | / |
| (iii) autobiographical memory cues | 2.88 (.46) | 4.75 (.30) | / | / |
| (iv) autobiographical memory cues | 2.87 (.50) | 4.72 (.39) | / | / |
| (v) autobiographical memory cues | 2.86 (.50) | 4.74 (.35) | / | / |
| (i) sentences - comprehension | 3.21 (.22) | 2.70 (.25) | 4.25 (1.13) | 6.18 (.34) |
| (ii) sentences - comprehension | 3.18 (.20) | 2.74 (.22) | 4.32 (1.22) | 6.10 (.48) |
| (iii) sentences - recall | 3.20 (.15) | 2.77 (.26) | 4.03 (0.99) | 6.07 (.46) |
| (iv) sentences - recall | 3.20 (.20) | 2.72 (.19) | 4.31 (1.27) | 6.13 (.46) |
